# Cortical hyperexcitability drives dying forward ALS symptoms and pathology in mice

**DOI:** 10.1101/2021.08.13.456320

**Authors:** Mouna Haidar, Aida Viden, Brittany Cuic, Taide Wang, Marius Rosier, Doris Tomas, Samuel A. Mills, Alistair Govier-Cole, Elvan Djouma, Sophia Luikinga, Valeria Rytova, Samantha K. Barton, David G. Gonsalvez, Lucy M. Palmer, Catriona McLean, Matthew C. Kiernan, Steve Vucic, Bradley J. Turner

**Author notes:** Corresponding author: Bradley Turner, Florey Institute of Neuroscience and Mental Health University of Melbourne, 30 Royal Parade, Parkville, VIC, Australia, 3052.

## Abstract

Amyotrophic lateral sclerosis (ALS) is a progressive fatal disorder caused by degeneration of motor neurons in the cortex and spinal cord. The origin of ALS in the central nervous system is unclear, however cortical hyperexcitability appears as an early and intrinsic feature of ALS and has been linked to degeneration of spinal motor neurons via a dying-forward mechanism. Here, we implement chemogenetics to validate the dying forward hypothesis of ALS in mice. We show that chronic hyperexcitability of corticomotoneurons induced by excitatory chemogenetics results in motor symptoms and core neuropathological hallmarks of ALS, including corticomotoneuron loss, corticospinal tract degeneration and reactive gliosis. Importantly, corticomotoneuron loss was sufficient to drive degeneration of spinal motor neurons and neuromuscular junctions (NMJs), associated with cytoplasmic TAR DNA binding protein 43 (TDP-43) pathology. These findings establish a cortical origin of ALS mediated by neuronal hyperexcitability, consistent with a dying forward mechanism of neurodegeneration.

## Introduction

Amyotrophic lateral sclerosis (ALS) is a progressive and universally fatal disorder caused by the selective degeneration of motor neurons in the brain and spinal cord^1^. Motor neurons are classified into upper populations in the motor cortex (corticomotoneurons) which synapse with lower populations in the spinal cord (spinal motor neurons) which innervate skeletal muscle by neuromuscular junctions (NMJs). The spatial and temporal relationship between cortical and spinal motor neuron loss in ALS remains unclear. Cortical hyperexcitability appears as an early and intrinsic feature of sporadic and familial ALS^2-5^, reflecting increased excitability of corticomotoneurons, a process that has been linked to degeneration of spinal motor neurons via trans-synaptic glutamate-mediated excitoxicity. These considerations represent a central premise of the ‘dying forward’ hypothesis of ALS which posits that spinal motor neuron loss is secondary to corticomotoneuron dysfunction^6^. Evidence for a potential cortical origin of ALS has been obtained using neuropathological and neurophysiological studies in patients, but to date, has been lacking in experimental models.

## Results

To induce cortical hyperexcitability in a spatiotemporally controlled manner in mice, we implemented a chemogenetic approach employing Designer Receptors Activated by Designer Drugs (DREADDs)^7,8^. We used viral delivery of the human modified muscarinic type 3 DREADD (hM3Dq) driven by the calcium/calmodulin dependent protein kinase II-α (CAMKIIα) promoter (AAV5-CAMKIIα-hM3Dq-mCherry) to target glutamatergic corticomotoneurons (Fig 1a, top panel). The excitatory hM3Dq DREADD was bilaterally injected and expressed in Layer V motor cortex in adult mice (Fig. 1a, middle panel). After establishing hM3Dq expression for 4-weeks, selective and chronic *in vivo* activation of corticomotoneurons was achieved by daily injections of the pharmacologically inert and highly potent designer drug clozapine-N-oxide (CNO) (5 mg/kg, IP) for 6 months (Fig 1a, bottom panel). Control hM3Dq mice were treated with saline vehicle. Negative control mice expressing fluorescent protein only without hM3Dq were treated with saline or CNO. Targeted expression of hM3Dq in Layer V of motor cortex in mice was achieved for all mouse cohorts and expression was mapped from hM3Dq-CNO mice from 2 cohorts and sex (Fig. 1b and Extended Data Fig. 1). Activation of corticomotoneurons was confirmed by increased c-Fos expression within CTIP2+ and hM3Dq+ corticomotor neurons, following CNO treatment in hM3Dq mice (hereafter hM3Dq-CNO), unlike hM3Dq mice treated with saline (hereafter hM3Dq-saline) and control mice expressing fluorescent protein treated with saline (hereafter Control-saline) or CNO (hereafter Control-CNO) (Fig. 1c). To confirm hM3Dq activity, we measured the effect of CNO on the intrinsic electrophysiological properties of hM3Dq+ corticomotoneurons with *ex vivo* patch clamp recordings. Bath application of 1 µM CNO induced increased firing rate (Fig. 1d, e) and increased membrane resistance (Fig. 1f, g), along with a reduction in rheobase (minimum current needed to induce action potential firing; Fig. 1h), compared to artificial cerebrospinal fluid (aCSF). hM3Dq+ corticomotoneuron firing was also shown with 5 µM CNO (Extended Data Fig. 2). Collectively, these data demonstrate the evolution of hyperexcitability and increased neuronal activation of corticomotoneurons expressing hM3Dq activated by CNO. Next, we examined whether chronic activation of corticomotoneurons can drive the development of ALS-related motor symptoms in mice. hM3Dq-CNO mice developed a distinct hindlimb clasping phenotype following 14 weeks of daily CNO treatment indicative of upper motor neuron signs, compared to hM3Dq-saline mice (Fig. 1i, Supplementary Video 1, Extended Data Fig. 3a). hM3Dq-CNO mice developed motor deficits shown by the rotarod test (Fig. 1j) and impaired hindlimb grip strength (Fig. 1k), compared to hM3Dq-saline mice. Control mice expressing fluorescent protein only treated with saline or CNO did not show deficits in these tests (Fig. 1 l, m). Spontaneous locomotor activity and weight gain were not affected by CNO treatment in any groups (Extended Data Fig. 3b, c), demonstrating a lack of off-target effects of CNO. We therefore subsequently focused on neuropathology in the hM3Dq-CNO and hM3Dq-saline groups. Thus, chronic hyperexcitability of corticomotoneurons drives key ALS-like motor symptoms in mice.

**Figure 1.**
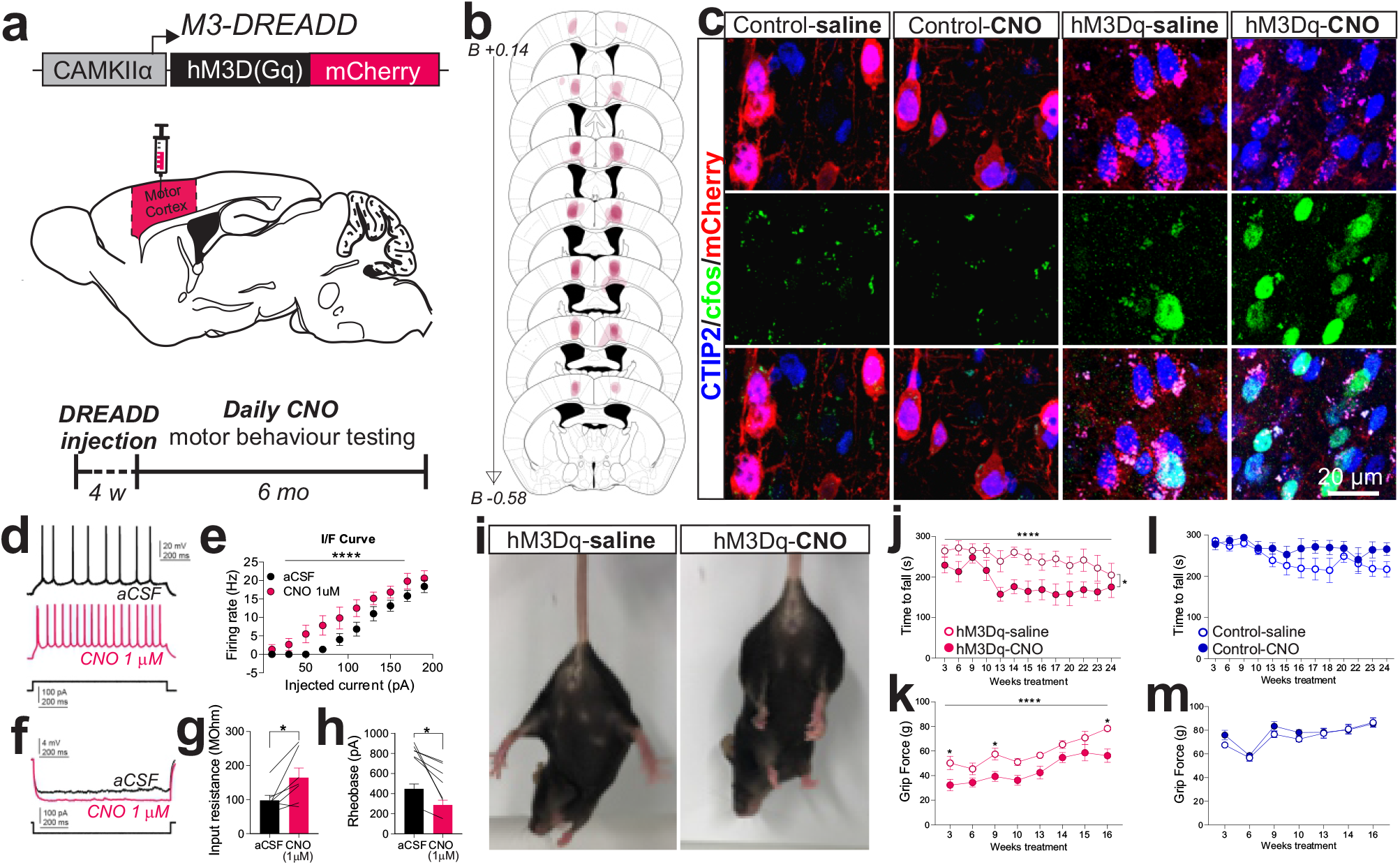
Hyperexcitability of corticomotoneurons conferred by hM3Dq activation induces ALS-like motor deficits in mice. **a**, Schematic of hM3Dq viral construct, stereotaxic injection into mouse primary motor cortex (M1), and outline of experimental procedure. **b**, Superimposed coronal sections depicting anatomically characterised hM3Dq+ neurons from each hM3Dq-CNO mice included in analysis [coronal brain images adapted from mouse brain atlas, Paxinos and Franklin 2001]. **c**, Representative images showing hM3Dq expression is confined to CTIP2+ (blue) neurons and c-Fos (green) levels are elevated in hM3Dq/mCherry+ neurons in hM3Dq mice treated with clozapine-N-oxide (CNO) relative to saline treated hM3Dq and saline and CNO treated control mice. **d**, Representative traces of the firing of a hM3Dq neuron before (aCSF, black) and after (CNO 1 µM, magenta) bath application of 1 µM CNO, in response to a 100 pA current step injected for 1,200 ms. **e**, Firing rates as a function of injected current of hM3Dq neurons. Bath application of 1 µM CNO increased the firing rate of hM3Dq neurons compared to pre-CNO application (two-way ANOVA, \*\*\*\**p* < 0.0001, *n* = 6 neurons). **f**, Representative traces of the voltage response of a hM3Dq neuron to a -90 pA current step injected for 1200 ms, before (aCSF, black) and after (CNO 1 µM, magenta) bath application of 1 µM CNO. **g**, Bath application of 1 µM CNO increased the membrane resistance of hM3Dq neurons compared to pre-CNO application (Wilcoxon paired test, \**p* = 0.0313, *n* = 7 neurons). **h**, Bath application of 1 µM CNO reduced the rheobase of hM3Dq neurons compared to pre-CNO application (Wilcoxon paired test, *p* = 0.0156, *n* = 7 neurons). **i**, Representative images of hindlimb splay in saline- and CNO-treated hM3Dq-expressing mice. Chronic CNO administration in hM3Dq-expressing mice (n=7) results in (**j**) impaired rotarod performance and (**k)** reduced hindlimb grip strength, compared to saline treated hM3Dq expressing mice (*n*=6). Control mice treated with saline (*n*=7) or CNO (*n*=8) show normal **l**, performance on a Rotarod and **m**, hindlimb grip strength. Data are expressed as mean ± SEM. **j – m**, Data were compared by two-way repeated measures ANOVA, \**p* < 0.05, **** *p* < 0.0001, compared with hM3Dq-saline groups.

To determine whether hyperexcitability of corticomotoneurons is sufficient to trigger cortical pathology, the total number of CTIP2+ neurons in Layer V of the motor cortex was assessed. Approximately 50% of CTIP2+ corticomotoneurons were significantly lost at 6 months post-treatment in hM3Dq-CNO mice, compared to hM3Dq-saline mice (Fig. 2a-b). In parallel with the loss of corticomotoneurons, significant degeneration of the corticospinal tract (CST) in the lumbar spinal cord occurred in hM3Dq-CNO mice, compared to hM3Dq-saline mice (Fig. 2c-d). Lastly, synaptophysin+ synapses with ChAT+ spinal motor neurons were significantly reduced in hM3Dq-CNO mice (Fig. 2e-g), demonstrating disruption of corticomotoneuron input into the anterior horn. Thus, cortical hyperexcitability drives degeneration of corticomotoneuron cell bodies, axons and projections to the spinal cord. Next, we used Spectral Confocal Reflectance Microscopy (SCoRe), a label-free structural imaging technique^9,10^, to evaluate compact myelin coverage of hM3Dq+ expressing axons arising from Layer V corticomotoneurons (Extended Data Fig. 4a). We observed no difference in SCoRe signal colocalised along axons of hM3Dq+ expressing corticomotoneurons (Extended Data Fig. 4a). In addition, we observed no difference in total SCoRe signal within dorsal funiculus of the lumbar spinal cord (Extended Data Fig. 4b). These data indicate that the hyperexcitability induced ALS-like motor symptoms and neuropathology arises primarily from corticomotor neurons, rather than a secondary effect of demyelination.

**Figure 2.**
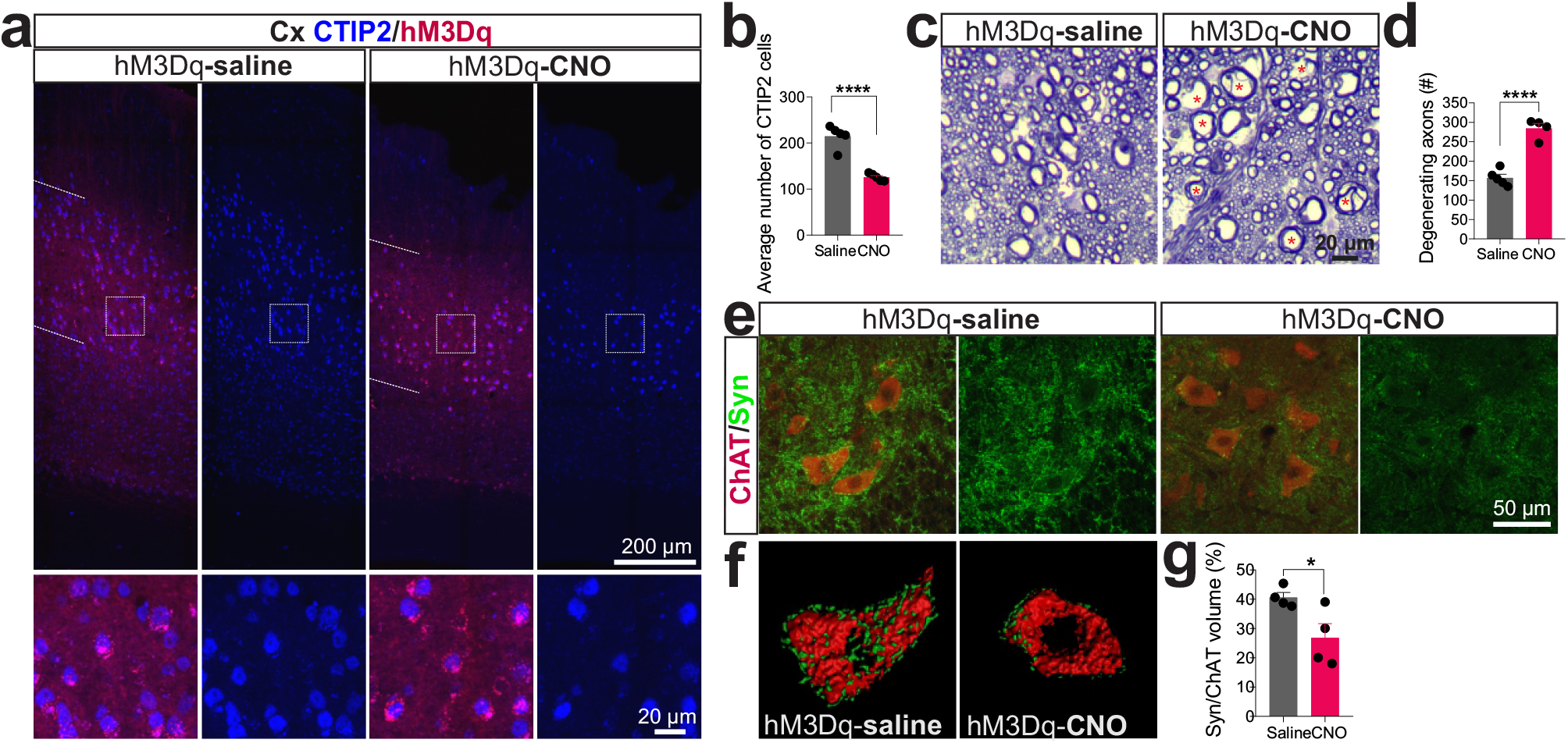
Cortical hyperexcitability drives corticomotoneuron loss and corticospinal tract axonal degeneration. **a**, Immunofluorescence images revealing widespread loss of CTIP2+ corticomotoneurons within the injection site of hM3Dq-CNO mice, compared to hM3Dq-saline mice. Boxed areas in **a** indicate the regions illustrated in magnified images showing selective loss of CTIP2-expressing corticomotoneurons in hM3Dq-CNO mice, compared to hM3Dq-saline controls. **b**, Quantitative analysis of average number of CTIP2+ cells in hM3Dq-saline (*n*=5) and hM3Dq-CNO (*n*=5) mice. **c**, Representative toluidine blue-stained sections of dorsal column from hM3Dq-saline and hM3Dq-CNO mice showing degenerating axons in hM3Dq-CNO mice (red asterisks). **d**, Quantitative analysis of average number of degenerating axons in hM3Dq-saline (*n*=5) and hM3Dq-CNO (*n*=4) mice. **e**, Representative confocal microscopy images of synaptophysin (green) and ChAT (red) in ventral horn of lumbar spinal cord and **f**, 3D rendering of spinal motor neurons indicated by arrowheads obtained by IMARIS software. **g**, Quantification analysis revealing a significant reduction in the proportion of synaptic contacts onto spinal motor neurons in hM3Dq-CNO mice (*n*=4), compared to hM3Dq-saline mice (*n*=4). Data are expressed as mean ± SEM. All data were compared by unpaired t-tests, \**p* < 0.05, \*\*\*\**p* < 0.0001 compared with hM3Dq-saline group.

To establish whether the advent of cortical hyperexcitability could drive pathology at the level of spinal motor neurons, we comprehensively analysed their cell bodies and peripheral synapses. Spinal motor neuron counts were similar at 6-months in hM3Dq-CNO and hM3Dq-saline mice (Fig. 3a-b). However, significant spinal motor neuron loss occurred in hM3Dq-CNO mice at 12-months post-CNO treatment, revealing a progression of pathology. We then examined NMJ morphology and denervation in hM3Dq-CNO and hM3Dq-saline mice at 6- and 12-months. NMJ areas were significantly reduced in gastrocnemius muscles of hM3Dq-CNO mice at both 6- and 12-months, compared to respective hM3Dq-saline controls (Fig. 3d, e). While NMJs were intact at 6-months in hM3Dq-CNO mice (Fig. 3d, f), 60% of NMJs were denervated in hM3Dq-CNO mice at 12-months, compared to hM3Dq-saline mice (Fig. 3e, f). These findings of spinal motor neuron degeneration induced by cortical hyperexcitability support an anterograde spreading mechanism of neurodegeneration in ALS.

**Figure 3.**
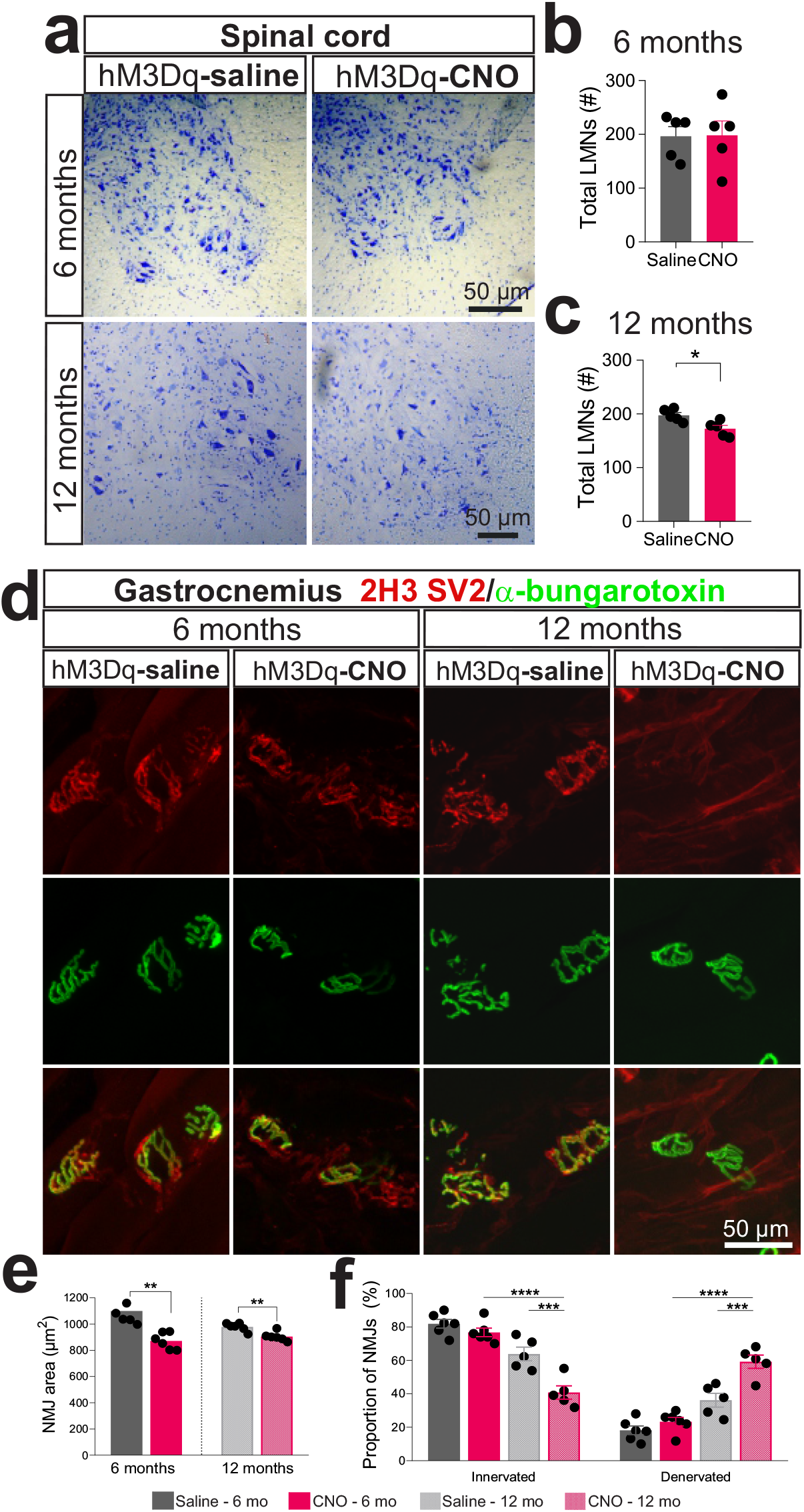
Hyperexcitability of corticomotoneurons drives spinal motor neuron loss and muscle denervation in mice. **a**, Representative images of spinal cord sections stained for Nissl from hM3Dq-saline and hM3Dq-CNO mice treated for 6 or 12 months. Quantification analysis of spinal motor neuron counts at (b) 6 months and (c) 12 months revealed significant loss in hM3Dq-CNO mice, relative to hM3Dq-saline mice at latter age. (Unpaired t-test, 6 months, *p* > 0.05; 12 months \**p* < 0.05, *n* = 5 per group). **d**, Representative confocal microscopy stacks of α-bungarotoxin (green) and 2H3/SV2 (red) in gastrocnemius of 6- and 12-month treated hM3Dq-saline and hM3Dq-CNO mice. **e**, Neuromuscular junction (NMJ) area is smaller in hM3Dq-CNO mice, compared to hM3Dq-saline mice following 6 and 12 months of treatment. (Unpaired t-test, 6 months, ** *p* < 0.01; 12 months, ** *p* < 0.01, *n* = 5-6 per group). **f**, NMJ denervation is observed following 12 months of treatment in hM3Dq-CNO mice, compared to hM3Dq-saline treated mice (Two-way ANOVA, *** *p* < 0.001; **** *p* < 0.0001, *n* = 5 per group). All data are expressed as mean ± SEM.

To assess if cortical hyperexcitability drives reactive gliosis characteristic of ALS, glial cell activation was studied at both cortical and spinal cord levels. Ionized calcium binding adaptor molecule 1 (Iba1+) microglial activation was present in Layer V of the motor cortex of hM3Dq-CNO mice (Fig. 4a), evidenced by reduced cell length, volume, retraction of microglial branches and terminals (Fig. 4b), unlike hM3Dq-saline mice. Here, we support the notion that microglial phenotypes are modulated by neuronal activity^11^, and show microglial activation is driven by corticomotoneuron hyperexcitability occurring in ALS. Importantly, activation of CD11b+ microglia also occurred in ventral horns of spinal cord in hM3Dq-CNO mice, compared to hM3Dq-saline mice at 6 months (Fig. 4c, d). Furthermore, GFAP+ astrocyte activation, indicated by reduced cell area, branches and processes, was shown in ventral horns of hM3Dq-CNO mice, compared to the hM3Dq-saline group (Fig. 4e, f). Hence, cortical hyperexcitability can propagate neuroinflammation to the spinal cord.

**Figure 4.**
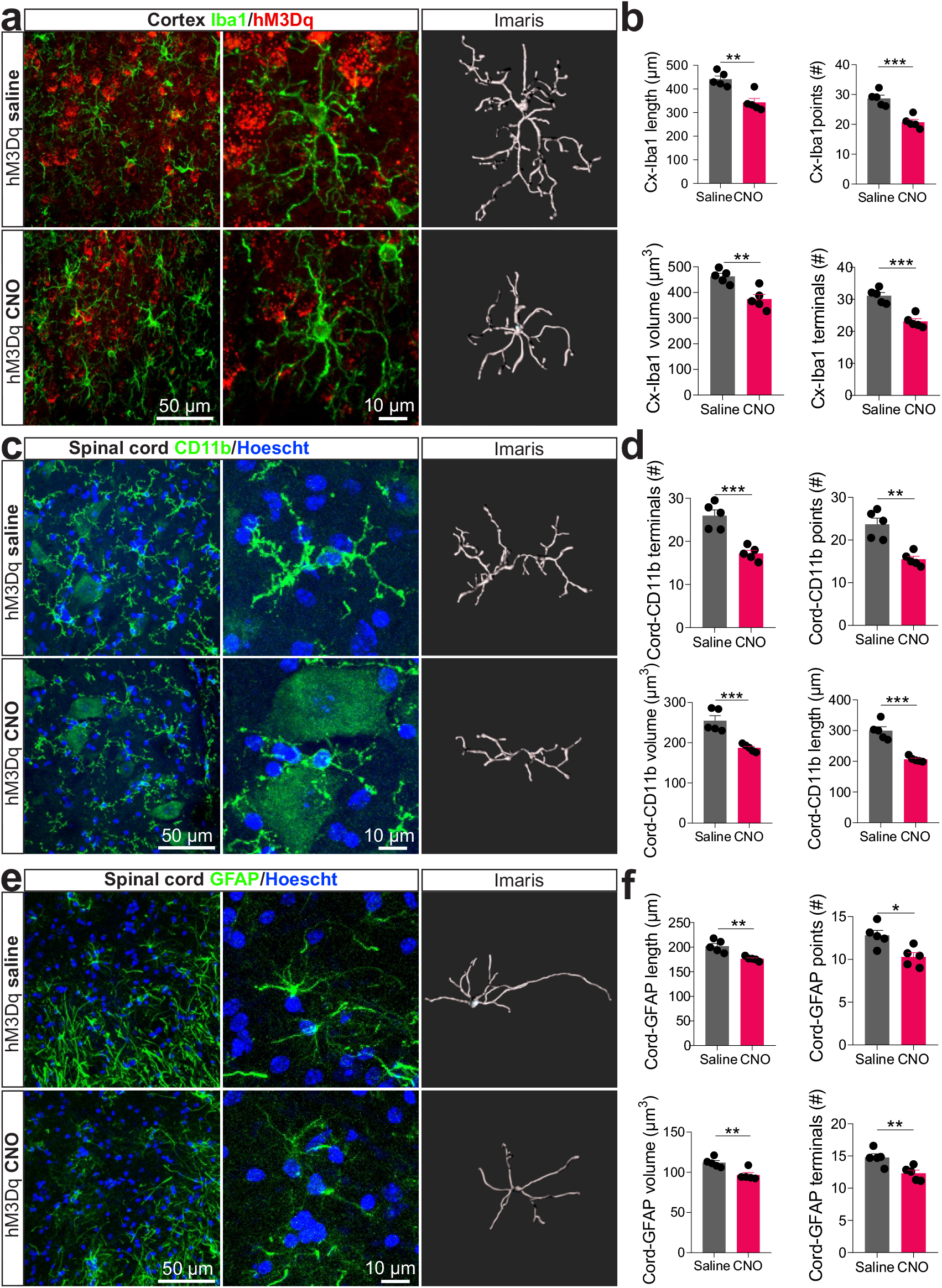
Corticomotoneuron hyperexcitability triggers reactive gliosis in the motor cortex and spinal cord. **a**, Representative confocal microscopy images of Iba1 (green) and hM3Dq-mCherry (red) within the injection site of motor cortex and 3D reconstruction of an individual Iba+ cell using filament extension in IMARIS. **b**, Overall signs of retracted Iba1 branches was observed in Iba1 branch length, branch volume, branch points and terminals. **c**, Representative confocal microscopy images of CD11b (green) and Hoechst (blue) within the ventral horn of spinal cord and 3D reconstruction of an individual CD11b+ cell using filament extension in IMARIS. **d**, Overall signs of retracted CD11b branches was observed in CD11b branch length, branch volume, branch points and terminals. **e**, Representative confocal microscopy images of GFAP (green) and Hoechst (blue) within the ventral horn of spinal cord and 3D reconstruction of an individual GFAP+ cell using filament extension in Imaris. **f**, Overall signs of retracted GFAP branches was observed in GFAP branch length, branch volume, branch points and terminals. Data are expressed as mean ± SEM. All data were compared by unpaired t-tests. \**p* < 0.05, \*\**p* < 0.01, \*\*\**p* < 0.001, *n* = 5 per group.

TDP-43 pathology as demonstrated by cytoplasmic accumulation and post-translational modifications remains the key hallmark feature of affected neurons in ALS^12^. To investigate TDP-43 pathology in our model, we used sequential biochemical fractionation of brain and spinal cord tissues from mice at 6 months. Phospho-TDP-43 (p-TDP-43) levels were significantly elevated in insoluble fractions of the frontal cortex in hM3Dq-CNO mice, compared to hM3Dq-saline animals (Fig. 5a, b). Insoluble total TDP-43 accumulation also occurred in the frontal cortex of hM3Dq-CNO mice, unlike hM3Dq-saline mice (Fig. 5a, c). There was a corresponding reduction of total TDP-43 levels in the soluble fraction of frontal cortex from hM3Dq-CNO mice (Extended Data Fig. 5a-b). Importantly, pTDP-43 levels were significantly higher in spinal cords of hM3Dq-CNO mice, compared to hM3Dq-saline mice (Fig. 5d, e). We next validated our biochemical findings with pTDP-43 immunohistochemistry in layer V of motor cortex and ventral horns of spinal cord. In contrast to hM3Dq-saline mice, corticomotoneurons in hM3Dq-CNO mice showed abundant cytoplasmic pTDP-43 accumulation and aggregation (Fig. 5g, top row). In addition, spinal motor neurons showed striking pTDP-43 accumulation and aggregation in the cytoplasm of hM3Dq-CNO mice, unlike hM3Dq-saline mice (Fig. 5g, bottom row). These results suggest that cortical hyperexcitability drives TDP-43 pathology in both motor cortex and spinal cord. Indeed, a recent study demonstrated that neuronal hyperexcitability drives cytoplasmic accumulation of TDP-43 in human patient iPSC-derived neurons^13^, in accordance with our findings in mice. Thus, we propose that TDP-43 pathology is a downstream event of neuronal hyperexcitability in ALS.

**Figure 5.**
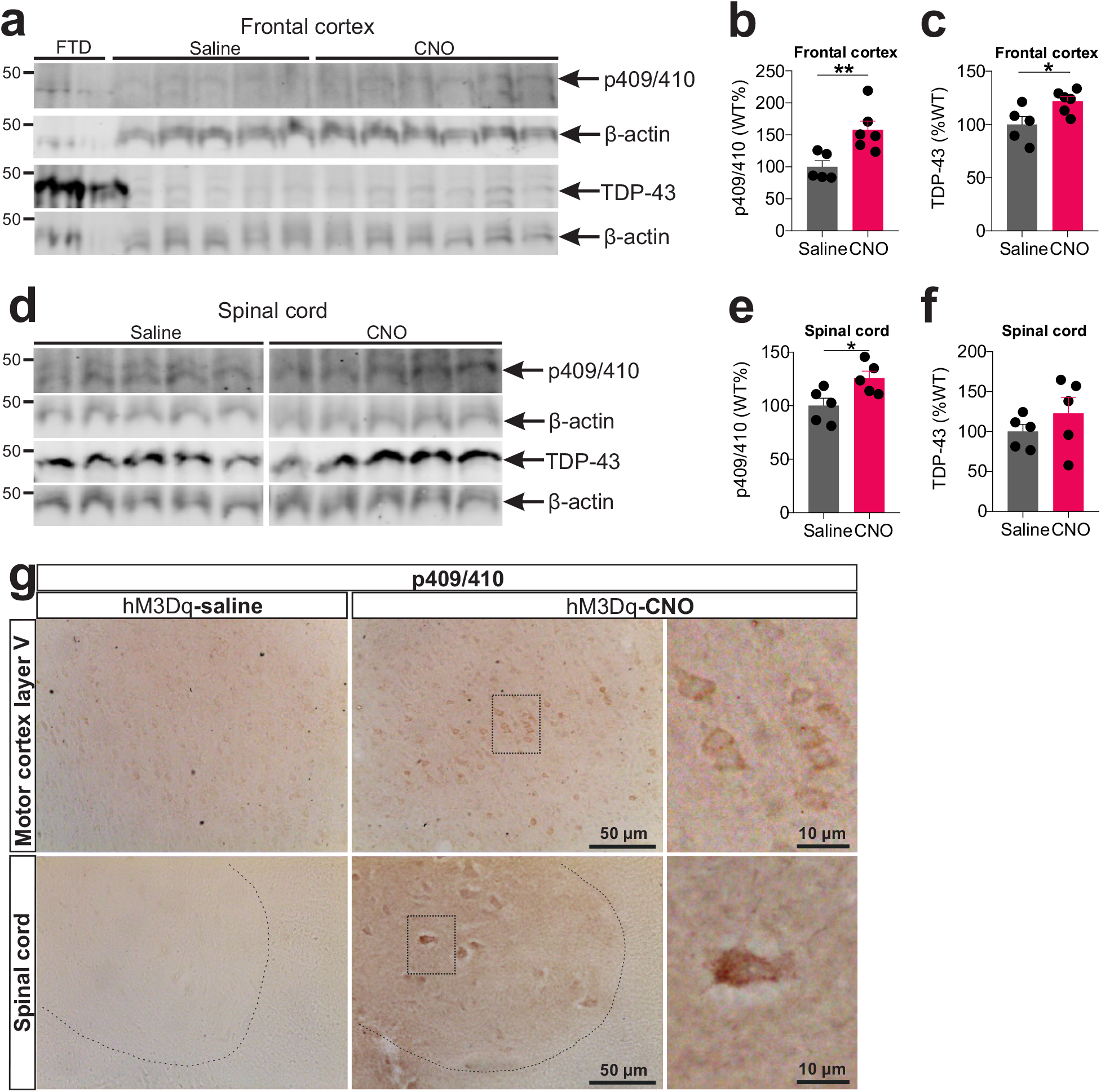
Hyperexcitability of corticomotoneurons drives TDP-43 pathology in motor cortex and spinal cord in mice. Immunoblot analysis of RIPA-insoluble phospho (p409/410)- and total TDP-43 in (**a**) frontal cortex and (**d**) lumbar spinal cord of hM3Dq-saline and hM3Dq-CNO mice, relative to β-actin levels. Quantification analysis revealing significantly higher levels of **b**, phospho- and **c**, total TDP-43 levels in frontal cortex of hM3Dq-CNO mice, compared to hM3Dq-saline mice. Significantly higher levels of phospho-TDP-43 are observed in **e**, lumbar spinal cord of hM3Dq-CNO mice, compared to hM3Dq-saline mice. Total TDP-43 levels in **f**, lumbar spinal cord in groups. Gel source data can be found in Supplementary Fig. 1. **g**, Representative immunohistochemistry images revealing the presence of cytoplasmic phospho-TDP-43 accumulation and aggregation in Layer V corticomotoneurons in the primary motor cortex (top row) and spinal motor neurons (bottom row) in hM3Dq-CNO mice, relative to hM3Dq-saline treated mice. Boxed areas (middle panel) indicate the magnified images (right panels). Data are expressed as mean ± SEM. All data were compared by unpaired t-tests, \**p* < 0.05, \*\**p* < 0.01, *n* = 5-6 per group.

## Discussion

It has been proposed that neurodegeneration in ALS may develop through a dying-forward mechanism originating in the motor cortex^6^, with cortical hyperexcitability the pathophysiological mechanism driving this process^2^. Using chemogenetics in mice, we demonstrate that cortical hyperexcitability invoked by excitatory DREADD activation induces not only corticomotoneuron degeneration, but loss of spinal motor neurons and NMJs, consistent with a dying forward mechanism of ALS onset and propagation. In our model, hyperexcitability of corticomotoneurons drives essential features of ALS, including age-dependent muscle weakness, corticomotoneuron and spinal motor neuron degeneration, reactive gliosis and importantly, TDP-43 pathology in affected motor neurons. We therefore conclude that cortical hyperexcitability is sufficient to drive ALS-like symptoms and pathology in mice.

Cortical hyperexcitability in ALS has been attributed to degeneration of inhibitory GABA-ergic circuits^14^ combined with hyperactivity of glutaminergic excitatory interneurons in the motor cortex^15^, with the net effect of causing overactivation of corticomotoneurons. We confined excitatory hM3Dq expression and activation to glutamatergic corticomotoneurons in Layer V of motor cortex, allowing us to precisely dissect the contribution of corticomotoneuron hyperexcitability to the pathological cascade of ALS in mice. G protein signalling through activation of hM3Dq recruits phospholipase C (PLC) to produce inositol phosphate (IP3) and diacylglycerol (DAG), leading to increased levels of calcium^16^. Interestingly, dysregulation of the PLC/IP3/DAG signalling pathway is linked to excitotoxicity in ALS^17^,^18^. Thus, our chemogenetic approach stimulates a signalling pathway leading to excitotoxicity occurring in ALS.

There are two major findings from this study. First, cortical hyperexcitability causes corticomotoneuron degeneration with secondary spinal motor neuron loss in our hM3Dq-CNO model, strongly supporting a dying forward proposal in ALS. Corticomotoneurons are glutamatergic and are thus likely to damage spinal motor neurons by glutamate-mediated excitotoxicity. The associated reactive microgliosis and activation of astrocytes in spinal cords of hM3Dq-CNO mice are also consistent with excitotoxicity. We also provide the first evidence that hyperexcitability of corticomotoneurons can trigger disruption of NMJs and muscle weakness which are hallmarks of lower motor neuron damage. Specifically, the processes of neurodegeneration as outlined in the present study are sufficient to cross and involve both compartments of the nervous system, central and peripheral, the key diagnostic feature of ALS ^19^. Secondly, and most importantly, this study demonstrates that corticomotoneuron hyperexcitability can drive the appearance of TDP-43 pathology, the pathological hallmark of ALS, placing neuronal excitability upstream of TDP-43 proteinopathy.

Altogether, our findings using chemogenetics in mice provide support for a cortical origin of ALS. Specifically, these results emphasise the primacy of corticomotoneuron dysfunction not only in ALS onset, but TDP-43 aggregation, reactive gliosis and muscle denervation which constitute key pathologies of ALS. Future studies will focus on harnessing chemogenetic approaches to mitigate cortical hyperexcitability as a potential intervention for ALS.

## Methods

### Mice

All experiments were performed with approval from The Florey Institute of Neuroscience and Mental Health Animal Ethics Committee (AEC, number: 17-097 and 20-159) and in accordance with the Australian National Health and Medical Research Council of Australia. Wild-type (WT) adult female and male mice on a C57BL/6J background were purchased from Animal Resources Centre, Western Australia, Australia. Mice were group housed in individually ventilated cages (IVC) under standard 12 h light-dark conditions (lights on 0700-1900) at 22°C with regular chow and water available *ad libitum*. Mice were randomly assigned to either experimental or control groups. Most behavioural experiments were performed on female mice from one cohort, unless otherwise stated. Male and female mice from two cohorts were used for histological experiments. Mice were weighed weekly for the entire duration of experimentation.

### Designer Receptors Exclusively Activated by Designer Drugs (DREADDs)

Plasmid adeno-associated viral (pAAV) vector encoding the human Gq-coupled M3 muscarinic receptor (hM3Dq) fused with mCherry under the control of CAMKIIa promoter (pAAV CAMKIIa-hM3D(Gq)-mCherry, AAV5; ≥ 2 × 10^11^ viral genomes (vg)/mL) or pAAV with enhanced green fluorescent protein (eGFP) under the control of CAMKIIa promoter (pAAV-CAMKIIa-EGFP, AAV5; ≥ 2 × 10^12^ vg/mL). The CAMKIIa promoter enables selective transgene expression in pyramidal neurons^20^. pAAV-CAMKIIa-hM3D(Gq)-mCherry pAAV-CAMKIIa-EGFP were a gift from Bryan Roth (Addgene plasmid # 50476 and # 50469).

### Stereotaxic surgery

6-8-week-old mice were anaesthetised with 5% isoflurane then secured in a small animal stereotaxic frame (David Kopf Instruments, Tujunga, CA) with 1.5-2% isoflurane delivered through a nose cone. Mice were subcutaneously injected with meloxicam (20 mg/kg) for analgesia and eyes were moistened with lubricating eye ointment (Lacri-Lube, Allergen, NJ). Viral vectors were loaded into a 2 µl Hamilton syringe mounted on a stereotaxic frame. Viral vectors (100 nl each hemisphere) were targeted to the primary motor cortex (M1) to the following coordinates: -0.2 mm anterior-posterior and ±1.1 mm medial-lateral from Bregma and from pia -0.8 mm dorsal-ventral. Viral vectors (100 nL in each hemisphere) was injected at a rate of 20 nl/min. Mice were given ∼2-4 weeks to recover to allow for viral expression before commencement of CNO administration and experimentation.

### Clozapine-N-oxide (CNO) administration

CNO was obtained from Advanced Molecular Technologies (catalogue number 34233-69-7). Stock solution of CNO was dissolved in Dimethyl Sulfoxide (DMSO) diluted in sterile saline (0.9% NaCl) and injected intraperitoneally (i.p) at a concentration of 5 mg/kg/day for 6 months. From 6-12 months, CNO was diluted in drinking water and administered at a final concentration of 5 mg/kg/day beyond. CNO was made fresh every other day for i.p and drinking water experiments.

### Motor behaviour

#### Rotarod

Motor coordination was tested weekly on an accelerating rotarod (Mouse Rota-rod, 47600, Ugo Basile) as described by us previously^21^. Mice were tested from 3 weeks post treatment until study termination. Before testing, mice were trained with one steady session and two ramping sessions at 5-40 rpm over 5 minutes with 10 minute rest intervals in between each session. On testing days, latency to fall was recorded from 2 sessions at 5-40 rpm over 5 minutes with a 10 minute rest interval. The data shown are the average of 2 sessions on each testing day.

#### Hindlimb grip strength

Hindlimb muscle force was measured weekly using a grip strength test (BIO-G53, Bioseb). Mice were held in front of a horizontal bar to allow only the hindlimb paws to grasp the bar. Muscle force (g) was determined by gently pulling the mice until both paws released the bar. Three measurements were taken in succession and the average muscle force was recorded for analysis.

#### Analysis of hindlimb clasping

Mice were lifted from the tail for 10 seconds and were scored as displaying hindlimb clasping if they retracted one or both paws. A percentage of affected mice was recorded for analysis.

### *Ex vivo* whole cell recordings

For electrophysiology experiments, a separate cohort of mice injected bilaterally with pAAV CAMKIIa-hM3D(Gq)-mCherry in M1 were deeply anaesthetized with isoflurane (3-5 % in 0.75 L/minute O_2_). Brains were transferred and cut in an ice-cold, oxygenated solution containing (in mM): 110 choline chloride, 11.6 Na-ascorbate, 3.1 Na-pyruvate, 26 NaHCO_3_, 2.5 KCl, 1.25 NaH_2_PO_4_, 0.5 CaCl_2_, 7 MgCl_2_ and 10 D-Glucose (Sigma). Coronal slices of M1 (300-350 µm thick) were cut with a vibrating micro slicer (Leica Vibratome 1000S) and incubated in a solution containing (in mM): 125 NaCl, 3 KCl, 1.25 NaH_2_PO_4_, 25 NaHCO_3_, 1 CaCl_2_, 6 MgCl_2_ and 10 D-Glucose at 35 °C for 20 to 30 minutes, followed by incubation at room temperature for at least 30 minutes before recording. All solutions were continuously bubbled with 95% O_2_/5% CO_2_ (Carbogen). Slices were visualized using Differential Interference Contrast (DIC) microscopy, and whole-cell patch clamp somatic recordings were made from visually identified fluorescent Layer V pyramidal neurons using a 565 nm LED light source (Thorlabs) for mCherry visualization. During recording, slices were constantly perfused at ∼2 ml/minute with carbogen-bubbled artificial cerebral spinal fluid (aCSF) containing (in mM): 125 NaCl, 25 NaHCO_3_, 3 KCl, 1.25 NaH_2_PO_4_, 2 CaCl_2_, 1 MgCl_2_ and 25 D-Glucose (osmolarity 300-305 mOsm) maintained at 30-34 °C. Patch pipettes were pulled from borosilicate glass (Sutter Instruments), had open tip resistance of 4-6 MΩ and were filled with an intracellular solution containing (in mM):135 potassium gluconate, 70 KCl, 10 sodium phosphocreatine, 10 HEPES, 4 Mg-ATP, 0.3 Na2-GTP and 0.3% biocytin adjusted to pH 7.25 with KOH (osmolarity 290 mOsm). Recordings were amplified using up to two dual channel amplifiers (Multiclamp 700B, Molecular Devices), filtered at 2 kHz, digitized (50 kHz), and acquired through an ADC/DAC data acquisition unit (Digidata 1440A, Molecular Devices) by using pClamp 10.7 software (Molecular Devices). All recordings were made in current clamp, and liquid junction potential was not compensated for. Incremental depolarising current steps from +10 to +190 pA (20 pA steps) and lasting for 1200 ms were injected to produce action potential firing and generate the frequency vs current (I/F) curve. Membrane resistance was measured by injecting a -90 pA current step lasting for 1200 ms. Rheobase was measured by injecting incremental depolarising current steps of +5 pA lasting for 5 ms until the cell discharged a single action potential. CNO (1 µM or 5 µM) was added to the recording aCSF, and cell properties were measured again using the same protocols 10 minutes after CNO reached the slice. When needed, the membrane potential under CNO was brought at the same value as in the control condition through current injection.

### Histology and data analysis

#### Tissue collection and processing

Mice were anaesthetised with 5% isoflurane inhalation and then administered sodium pentobarbital (100 mg/kg, 0.1 ml, i.p) and transcardially perfused with 0.1 M phosphate buffer saline (PBS) followed by 4% paraformaldehyde (PFA) in 0.1 M phosphate buffer (PB). To validate neuronal activation for c-Fos expression, all mice were treated with saline or CNO and perfused 2 hours later. Mice were decapitated and brain and spinal cord were dissected and post-fixed in 4% PFA in 0.1 M PB for 1 hour at room temperature, then transferred to a solution of 30% sucrose in 0.1 M PBS and kept at 4°C until tissue sunk. Gastrocnemius muscle was dissected immediately after perfusion with 0.1 M PBS (prior to perfusion with PFA), and post-fixed in 4% PFA in 0.1 M PB for 10 minutes and washed in 0.1 M PBS three times then transferred to a solution of 30% sucrose in 0.1 M PBS and kept at 4°C until tissue sunk. Fixed brains were embedded on optimal cutting temperature (OCT, Tissue Tek) and snap-frozen on dry-ice and 40 µm coronal sections were cut and collected into 8 series spanning from +0.50 mm to -0.94 mm from Bregma using a cryostat at -18°C and stored in cryoprotectant solution (30% ethylene glycol, 30% glycerol, 0.05M PB) at -20°C. Lumbar region (L1-6) was dissected from fixed spinal cords and embedded in OCT and snap frozen on dry ice and 20 µm coronal sections were cut and mounted on SuperFrost^®^ Ultra Plus slides in 10 series, unless otherwise stated. Gastrocnemius was embedded in OCT and snap frozen on dry ice and 100 µm coronal sections were cut and mounted on SuperFrost^®^ Ultra Plus slides in 3 series. The investigators were blinded to the treatment groups during processing and data analysis for all experiments described below. All images for immunofluorescence experiments were captured on a Zeiss LSM 780 confocal laser scanning microscope (Carl Zeiss AG, Oberkochen, Germany), unless otherwise stated.

#### Spinal motor neuron quantification

For spinal motor neuron counts, slide mounted sections were stained with 0.5% cresyl violet with 0.04% acetic acid, dehydrated in ethanol, dipped in xylene and cover slipped with DePeX Mounting Media (BDH Chemicals). Images for quantification were acquired on a Zeiss Primo Star bright field microscope using a 10x/0.25 air objective lens. Alpha motor neurons were identified as large polygonal Nissl-positive neurons located within the ventral horn. Only cells with a clear nucleolus were counted to avoid double counting of neurons. 10 ventral horn sections per mouse were manually counted using event markers in Zen software (blue edition).

#### Corticospinal tract axon analysis

Perfusion fixed lumbar spinal cords were post-fixed in 2% osmium tetroxide, 1.5% potassium ferrocyanide in 0.05 M cacodylate buffer, washed, dehydrated and embedded in Spurrs Resin. 1 µm cross sections were cut and stained with 1% toluidine blue 2% sodium borate in distilled water. Images were acquired on a Zeiss Primo Star bright field microscope using a 40x/0.65 oil objective lens. 5 images were acquired and analysed from the dorsal funiculus from 3 sections per mouse and degenerating axons were manually counted using event markers in Zen software (blue edition). Degenerating axons were determined by decompaction of myelin structure, abnormal toluidine staining accumulation and axonal swelling.

#### Fos activation

Free-floating (40 μm) brain sections around the site of injection were washed three times and then blocked in 0.1 M PBS with 10% normal donkey serum and 0.5% Triton X-100 for 1 hour at room temperature. Sections were then incubated overnight in primary antibodies; rat anti-CTIP2 (1:500, ab18465, Abcam), rabbit anti-phospho-c-fos (1:200, 25348, Cell Signalling) and chicken anti-RFP (1:1000; 600-901-379S, Rockland). Tissue was then washed three times in 0.1 M PBS, blocked in 10% normal donkey serum and 0.5% Triton X-100 in 0.1 M PBS for 1 hour at room temperature and incubated in secondary antibodies; donkey anti-rabbit Alexa-Fluor 647 (1:400, 711-545-152, Jackson ImmunoResearch), donkey anti-rat Alexa-Fluor 488 (1:400, 712-605-153, Jackson ImmunoResearch) and donkey anti-chicken Alexa-Fluor 594 (1:400; 703-585-155, Jackson ImmunoResearch) for 2 hours at room temperature in 1% normal donkey serum and 0.1% Triton X-100 in 0.1 M PBS. Representative z stack images were captured using a 63x/1.40 oil objective lens with a 0.4 step in the z direction.

#### Corticomotoneuron quantification

To quantify corticomotoneurons, 3 brain sections processed for CTIP2 immunofluorescence (above) spanning from 0.62 mm to -0.70 mm from Bregma were chosen for analysis. Mosaic z stack images using a 20x/0.8 air objective lens images covering the entire M1 from each section were obtained with 10% overlap on the X-Y plane and 1 µm interval in the Z plane. M1 boundaries were identified by anatomical landmarks and outlined on each section for consistency of fields for analysis. Z-stacks were projected into a single image. The total number of CTIP2+ cells within the defined M1 field were manually counted using event markers in Zen software (blue edition). The data shown is the average number of CTIP2+ cells from each section and M1 counted per mouse.

#### Three-dimensional reconstruction of synaptic contacts on motor neurons and analysis

Slide mounted lumbar spinal cord sections (20 μm, L1-6) were incubated in an 1 mM ethylenediaminetetraacetic acid (EDTA) buffer solution (pH6) in dH_2_O at 90°C for 10 minutes, then briefly submerged in dH_2_O then washed in 0.1 M PBS and blocked. Tissue was then incubated 48 hours at 4°C in primary antibodies; goat anti-ChAT (1:250, AB144P, Merck Millipore) and rabbit anti-synaptophysin (1:500, 101-002, Synaptic Systems). Following washes and blocking (described above), tissue was incubated in secondary antibodies; donkey anti-goat Alexa-Fluor 594 (1:400, A11058, ThermoFisher) and donkey anti-rabbit Alexa-Fluor 488 (1:400, 711-545-152, Jackson ImmunoResearch). Z stack images were captured using a 40x/1.4 oil objective lens, 1.2 zoom with a 0.4μm step in the z direction. A three-dimensional reconstruction of synaptophysin+ synapses and ChAT+ cells was created using the Surface Wizard in Bitplane Imaris (version 8.4.1). To determine synaptic contacts onto alpha motor neurons the proportion of synaptophysin/ChAT volume was assessed from individual alpha motor neurons (> 20 µm in diameter) and the data shown is the average % of synaptophysin/ChAT from induvial motor neurons. ∼10-15 motor neurons were sampled per mouse.

#### Three-dimensional reconstruction of NMJ surface and analysis of innervation

Slide mounted gastrocnemius sections were washed three times in 0.1 M PBS, permeabilised for 30 minutes in 2% TritonX-100 in PBS then blocked in 10% normal donkey serum in 0.1 M PBS for 30 minutes at room temperature. Sections were then incubated at room temperature overnight in primary antibodies; SV2-5 (1:100; AB 2315387, Hybridoma b), mouse anti-2H3 (1:50; AB 2314897, Hybridoma b) in 0.1 M PBS. The following day, the tissue was washed three times and incubated in secondary antibody Alexa-Fluor 555 donkey anti-mouse (1:250, A-31570, ThermoFisher) and α-bungarotoxin Alexa Fluor 488 conjugate (1:1000; B13422, Invitrogen) in PBS for 2 hours. Images were captured using a 20x/0.8 air objective lens and taken with a 1 μm step in the z direction. Z-stacks were projected into a single image. For quantification of innervation status, BTX-labelled NMJs with clear overlap with presynaptic and postsynaptic labelling were defined as innervated. ∼ 80-100 NMJs were counted for innervation per mouse. To assess NMJ size, a three-dimensional reconstruction of each NMJ was created using the Surface Wizard in Bitplane Imaris (version 8.4.1). ∼ 50 NMJ surfaces were reconstructed per mouse.

#### Three-dimensional reconstruction of microglia and astrocytes and analysis

Free-floating (40 μm) brain sections around the site of injection were washed and blocked (described above). Antigen retrieval was performed on slide mounted lumbar (L1-6) spinal cord sections (20 μm; 90°C in EDTA for 10 minutes then rinsed with room temperature dH_2_O) prior to being washed and blocked. Brain sections were then incubated overnight at room temperature in primary antibodies; rabbit anti-Iba1 (1:1000; 019-19741, Wako), rat anti-GFAP (1:500; 13-0300, ThermoFisher) and chicken anti-RFP (1:1000; 600-901-379S, Rockland). Spinal cord sections were incubated for 48 hours at 4°C in primary antibodies; rabbit anti-CD11b (1:100; 133357, Abcam), mouse anti-GFAP (1:250; MAB360, Millipore) and goat anti-ChAT (1:250, ab144p, Abcam). Following washes and blocking (described above), tissue was incubated in the following secondary antibodies; donkey anti-rabbit Alexa-Flour 488 (1:400; 711-545-152, Jackson ImmunoResearch), donkey anti-chicken Alexa-Fluor 594 (1:400; 703-585-155, Jackson ImmunoResearch), donkey anti-goat Alexa-Fluor 594 (1:400; ab150132, Jackson ImmunoResearch), donkey anti-mouse Alexa-Fluor 647 (1:400; A31571, Thermo Fisher, Jackson ImmunoResearch) and donkey anti-rat Alexa-Fluor 647 (1:400; 712-605-153, Jackson ImmunoResearch) for 2 hours at room temperature. Images were captured using a 40x/1.4 oil objective lens. Z stack images were taken within the injection site with 0.4 μm step in the z direction. Using Imaris Filament Tracer Wizard, a three-dimensional surface was reconstructed for each individual microglia and astrocytes and analysed by the software. 10 microglia and astrocytes were reconstructed per mouse.

#### pTDP-43 immunohistochemistry

The tissue was washed 3 × 5 minutes in PBS, then endogenous peroxidase activity was blocked with 10% H_2_O_2_, 10% methanol and 80% PBS, at room temperature for 15 minutes. The sections were washed again 3 × 5 minutes in PBS and incubated overnight with rat anti-phospho-TDP primary antibody (1:100, 829901, BioLegend), in a dilution buffer (0.05% Tween, 2% NDS, in PBS). The following day, the tissue was washed 3 × 5 minutes in PBS prior to a 2 hour incubation with a biotin-conjugated donkey anti-rat secondary antibody (1:500, 712-065-153, Jackson ImmunoResearch). The primary–secondary antibody complexes were detected via peroxidase-driven precipitation of diaminobenzidine (DAB). This was carried out using avidin-biotin complex (ABC, 1 hour, Vectastain ABC Kit; Vector Laboratories) and then incubated with DAB (0.5 mg/mL for 5 minutes), which was precipitated by the addition of 1% (w/v) H_2_O_2_. The slides were then dehydrated in alcohol and xylene and cover slipped with DePeX Mounting Media (BDH Chemicals). Images were captured on a Leica DM6000 upright microscope using a 40x/0.65 air objective lens.

### Spectral Confocal Reflectance Microscopy (SCoRe)

Myelination of corticomotoneurons transfected with hM3Dq were assessed using SCoRe microscopy. Acquisition of the SCoRe signal and hM3Dq were performed within the same regions, in order to restrict analysis of the SCoRe signal to within the axons of transfected neurons. Anatomically this corresponded to cortical layers V and II/III. hM3Dq was imaged using standard confocal imaging by excitation of the reporter fluorophore, while myelin was imaged using reflected excitation laser light of wavelengths 488nm, 561nm and 647nm, as described previously^10^. Myelination within the descending corticospinal tract (CST) was assessed in transverse section at the spinal cord level L1. The CST was anatomically defined using the Atlas of the Mouse Spinal Cord^22^. All microscopy was performed on a Zeiss LSM980 AiryScan2 confocal microscope using a 40x/1.3 oil immersion objective lens at resolution determined by Nyquist sampling theorem. All imaging and subsequent image analysis was performed while blinded to the experimental groups. Image analysis was performed in FIJI. The three SCoRe channels (488 nm, 561 nm and 647 nm) were combined to create a single channel of comprehensive SCoRe signal. For cortical analysis, a median filter was applied and hM3Dq and SCoRe were manually thresholded to create binary masks. These masks were used to calculate 1) the area of hM3Dq positive axons in the field of view, and 2) the percentage of this area that was also positive for SCoRe signal. This allowed us to quantify the myelin coverage only in transfected cells. For CST analysis, images were segmented manually to isolate the CST, and the volume of CST within the z-stack was measured. The SCoRe signal was then manually thresholded and the volume of SCoRe was obtained and expressed as a density of total CST volume. After image analysis samples were unblinded and Students t-test performed.

### Protein extraction and quantification

Mice were anaesthetised with 5% isoflurane inhalation and then administered sodium pentobarbital (100 mg/kg, 0.1 mL, i.p) and transcardially perfused with 0.1 M PB to thoroughly remove blood and the frontal cortex and lumbar spinal cord were dissected and snap frozen and stored at −80°C. For a positive control, motor cortex tissue was used from a human patient with frontotemporal dementia, with protein extracted alongside the mouse tissue. Use of the human post-mortem tissue samples was approved by the Medicine and Dentistry Human Ethics Sub-Committee of the Human Research Ethics Committee at University of Melbourne; ethics #1852824. Tissues were thawed and sonicated in ice-cold RIPA buffer (50 mM Tris-Cl, pH 7.4, 150 mM NaCl, 1% (v/v) TX-100, 0.1% (w/v) SDS (AMRESCO, 0227), 1% (w/v) sodium deoxycholate (Sigma, D6750) with freshly added phosphatase inhibitors (50 mM NaF and 1 mM Na_3_VO_4_) and 1% (v/v) mammalian protease inhibitor cocktail (Sigma). Sonication was conducted at 50% output (Q55 Sonicator, Sonica, Newtown, CT, USA) with brief pulses applied over 5–10 s until tissue particulates were no longer visible. Samples were then stored on ice for 20 minutes and centrifuged at 21,000 g for 25 minutes at 4°C and the supernatant taken as the RIPA-soluble fraction. The pellet was then washed with RIPA buffer by sonication, and the supernatant was then discarded. This sequential pellet was then sonicated with 3x v/w urea buffer (7M urea, 2M thiourea, 4% CHAPS, 30mM Tris, pH 8.5), and centrifuged at 22 °C, 21,000g for 30 min. The supernatant was collected as the RIPA-insoluble/Urea-soluble fraction. RIPA soluble supernatants were quantified for protein concentration using the BCA assay according to the manufacturer’s protocol (Pierce® BCA assay kit, Thermo Fisher, 23225). RIPA soluble samples (30 μg) and urea soluble samples were denatured by boiling in Laemmli buffer containing 10% (v/v) β-mercaptoethanol.

### Immunoblotting

For both RIPA and Urea soluble fractions, samples were separated by electrophoresis on 4-20% Mini-PROTEAN® TGX Stain-Free™ gels (Bio-Rad Laboratories, NSW, Australia) in running buffer (0.1% (w/v) SDS, 14.8% (w/v) glycine in 100mM Tris-HCl, pH 8.2). Proteins were then transferred onto Transblot Turbo LF PVDF membrane (Bio-Rad) at 25V for 10 minutes using a TransBlot® Turbo™ Transfer System (Bio-Rad). Membranes were blocked with 5% BSA in TBST, and antibodies were diluted with SignalBoost™Immunoreaction Enhancer Kit (Merck). Primary antibodies included phospho-TDP-43 (Ser409/410) (Rabbit, 1:1000, 22309-1-AP, Proteintech) and TDP-43 (Rabbit, 1:1000,10782-2-AP, Proteintech). Blots were washed three times in TBST the following day in 10 minute intervals and incubated with 800CW secondary antibody (1:10,000, Li-Cor Biosciences), in addition to rhodamine anti-β-actin (1:5,000, Bio-Rad, 12004163) for 1 hour. Blots were washed three times in TBST in 10 minute intervals at room temperature. Membranes were imaged on a ChemiDoc™ MP (Bio-Rad). For analysis, blots were quantified using ImageJ software (Rasband WS, NIH, Bethesda, MD, http://rsb.info.nih.) by taking the mean grey value of bands for the target protein normalised to β-actin levels after subtracting background intensity. Results were expressed as a percentage of WT (100%). The investigator was blinded to the treatment group of the samples during the protocol and analysis.

## Acknowledgements

This work was supported by a FightMND IMPACT grant (M.H), MND Research Australia grant (M.H), Bethlehem Griffiths Research Foundation grant (M.H), University of Melbourne Early Career Research Grant (M.H) and Stafford Fox Medical Research Foundation grant (B.J.T). T.W. was supported by a FightMND Angie Cunningham PhD Scholarship and Grant, and B.J.T was supported by a NHMRC-ARC Dementia Research Leadership Fellowship 1137024 (B.J.T). S.K.B. was supported by a Rebecca L. Cooper Al & Val Rosenstrauss Medical Research Fellowship. We acknowledge the Centre for Advanced Histology and Microscopy at the Peter MacCallum Cancer Centre for the support of this work in sample preparation for corticospinal tract axon analysis. Human brain tissues were received from the Victorian Brain Bank, supported by The Florey, The Alfred, Victorian Institute of Forensic Medicine and Coroners Court of Victoria and funded in part by Parkinson’s Victoria, MND Victoria, FightMND, Yulgilbar Foundation and Ian and Maria Cootes. The Florey Institute of Neuroscience and Mental Health acknowledges support from the Victorian Government, in particular, funding from the Operational Infrastructure Support Grant.

## Author Contributions

M.H and B.J.T designed all experiments and wrote the paper. M.H and A.V performed stereotaxic surgery, administered daily CNO, assessed motor behaviour, perfused animals, processed tissue, performed immunofluorescence experiments, imaged, analysed the data, and statistical analysis. B.C and S.L assisted in CNO administration and motor behaviour. D.T assisted in stereotaxic surgery, mouse perfusions, tissue processing and motor behaviour. T.W performed western blot experiments and statistical analysis. M.R and L.M.P planned electrophysiology experiments. M.R performed electrophysiology experiments, analysed the data, and statistical analysis. S.A.M, A.G and D.G.G planned SCoRe experiments. A.G processed brain and spinal cord tissue for SCoRe and statistical analysis. S.A.M performed SCoRe imaging and analysed the data. V.R performed pTDP-43 immunohistochemistry and imaging and assisted with stereotaxic surgery. E.D assisted with CNO dosages and study design. S.K.B provided human tissue for western blot experiments and advised on analysis of axonal degeneration. C.M provided access to human brain tissues from the Victorian Brain Bank. S.V and M.C.K contributed to manuscript drafting and editing. All authors reviewed the manuscript prior to submission.

## Extended Data Figure Legends

**Extended Data Figure 1. Anatomical characterisation of hM3Dq expression in primary motor cortex. a**, Representative mosaic and higher magnification z stack mosaic revealing widespread bilateral expression of hM3Dq/mCherry+ neurons (red) within CTIP2+ neurons (a marker of Layer V cortical neurons). **b**, Superimposed coronal sections depicting anatomically characterised hM3Dq+ neurons from each sample from two independent cohorts and sex included in neuropathological analysis [coronal brain images adapted from mouse brain atlas, Paxinos and Franklin 2001].

**Extended Data Figure 2. Validation of corticomotoneuron hyperexcitability. a**, Representative micrographs depicting hM3Dq+ neuron and patch electrode under an infrared lamp (top) and fluorescent microscope (bottom, scale bar, 10 µm). **b**, Representative traces of the firing of a hM3Dq neuron before (black) and after (magenta) bath application of 5 µM CNO, in response to a 100 pA current step injected for 1,200 ms. **c**, Firing rates as a function of injected current of hM3Dq neurons. Bath application of 5 µM CNO increased the firing rate of hM3Dq neurons compared to pre-CNO application (two-way ANOVA, *p* = 0.0275, *n* = 5 neurons). **d**, Representative traces of the voltage response of a hM3Dq neuron to a -90 pA current step injected for 1,200 ms, before (black) and after (magenta) bath application of 5 µM CNO. **e**, Bath application of 5 µM CNO increased the membrane resistance of hM3Dq neurons, compared to pre-CNO application (Wilcoxon paired test, *p* = 0.0313, *n* = 7). **f**, Bath application of 5 µM CNO reduced the rheobase of hM3Dq neurons, compared to pre-CNO application (Wilcoxon paired test, *p* = 0.0156, *n* = 7 neurons).

**Extended Data Figure 3. Hindlimb clasping, spontaneous locomotor activity and body weight of mouse groups. a**, Percent of hM3Dq-saline (*n*=6) and hM3Dq-CNO mice (*n*=7) clasping from 14 to 18 weeks. **b**, Voluntary locomotor activity in a locomotor cell did not differ between hM3Dq (*n* = 10-12 per group) and control mice (*n* = 7-8 per group) following an acute treatment with clozapine-N-oxide (CNO). **c**, Body weights of hM3Dq (*n* = 10-11 per group) and control mice (*n* = 7-8 per group) treated with saline or CNO mice did not differ over time. Data are expressed as mean ± SEM. Data were compared by two-way repeated measures ANOVA, *p* > 0.05.

**Extended Data Figure 4. Myelin coverage is unaffected in motor cortex and corticospinal tract in hM3Dq-CNO mice. a**, Representative Spectral Confocal Reflectance (SCoRe) images of myelin signal (cyan) within the axons (red) of hM3Dq neurons, and within the dorsal funiculus of the lumbar spinal cord. **b**, Quantification of the myelinated area positive for SCoRe signal does not differ between hM3Dq-CNO and hM3Dq-saline mice in Layer V motor cortex (Cx), and corticospinal tract (CST) within the dorsal funiculus of lumbar spinal cord. Data are expressed as mean ± SEM. All data were compared by unpaired t-tests, *p* > 0.05, *n* = 5 per group.

**Extended Data Figure 5. Protein levels of RIPA-soluble TDP-43 in frontal cortex are significantly reduced in hM3Dq-CNO mice**. Immunoblot analysis of RIPA-soluble TDP-43 in frontal cortex **a**, and **c**, lumbar spinal cord of hM3Dq-saline and hM3Dq-CNO mice, relative to β-actin levels. **b**, Quantification analysis revealing significantly lower levels of TDP-43 levels in frontal cortex of hM3Dq-CNO mice, compared to hM3Dq-saline mice. **d**, TDP-43 levels in lumbar spinal cord in hM3Dq-CNO mice, compared to hM3Dq-saline mice. Gel source data can be found in Supplementary Fig. 1. Data are expressed as mean ± SEM. All data were compared by unpaired t-tests, * *p* < 0.05, *n* = 5-6 per group.

