## Extended Data for "Cortical hyperexcitability drives dying forward ALS symptoms and pathology in mice"

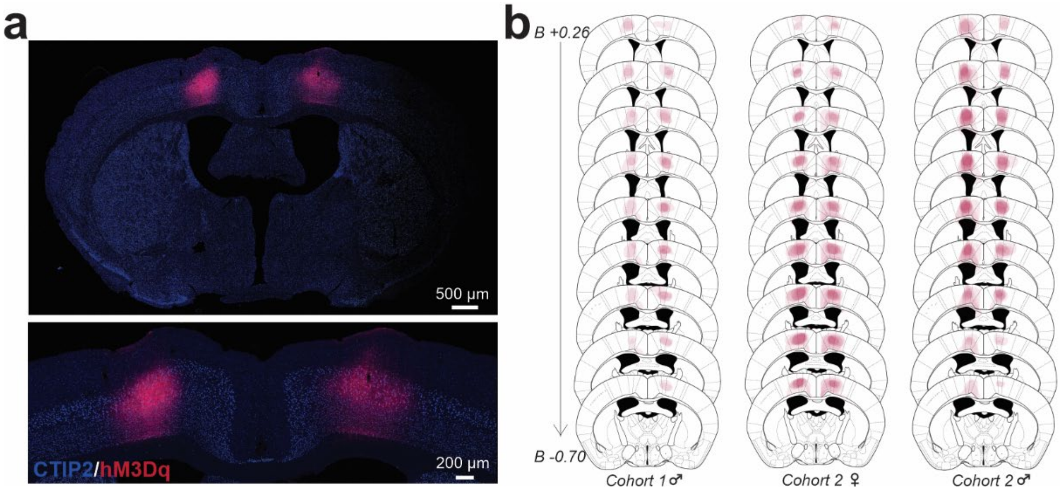


**Fig. S1. Anatomical characterisation of hM3Dq expression in primary motor cortex.** a, Representative mosaic and higher magnification z stack mosaic revealing widespread bilateral expression of hM3Dq/mCherry+ neurons (red) within CTIP2+ neurons (a marker of Layer V cortical neurons). b, Superimposed coronal sections depicting anatomically characterised hM3Dq+ neurons from each sample from two independent cohorts and sex included in neuropathological analysis [coronal brain images adapted from mouse brain atlas, Paxinos and Franklin 2001].

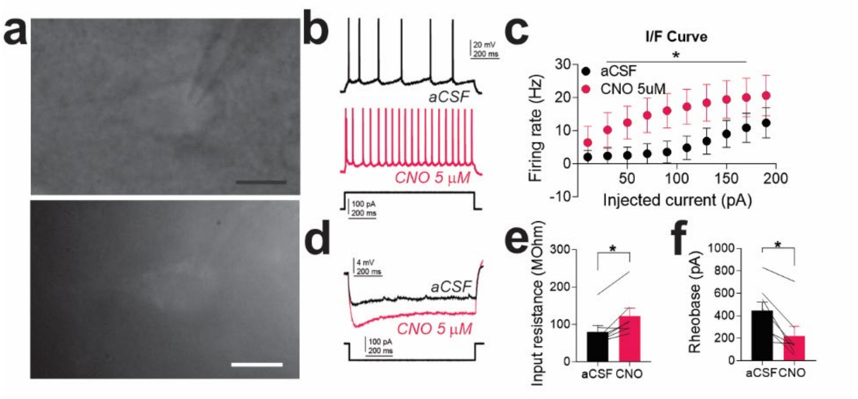


**Fig. S2. Validation of corticomotoneuron hyperexcitability.** a, Representative micrographs depicting hM3Dq+ neuron and patch electrode under an infrared lamp (top) and fluorescent microscope (bottom, scale bar, 10 µm). b, Representative traces of the firing of a hM3Dq neuron before (black) and after (magenta) bath application of 5 µM CNO, in response to a 100 pA current step injected for 1,200 ms. c, Firing rates as a function of injected current of hM3Dq neurons. Bath application of 5 µM CNO increased the firing rate of hM3Dq neurons compared to pre-CNO application (two-way ANOVA, $p = 0.0275$, $n = 5$ neurons). d, Representative traces of the voltage response of a hM3Dq neuron to a -90 pA current step injected for 1,200 ms, before (black) and after (magenta) bath application of 5 µM CNO. e, Bath application of 5 µM CNO increased the membrane resistance of hM3Dq neurons, compared to pre-CNO application (Wilcoxon paired test, $p = 0.0313$, $n = 7$). f, Bath application of 5 µM CNO reduced the rheobase of hM3Dq neurons, compared to pre-CNO application (Wilcoxon paired test, $p = 0.0156$, $n = 7$ neurons).

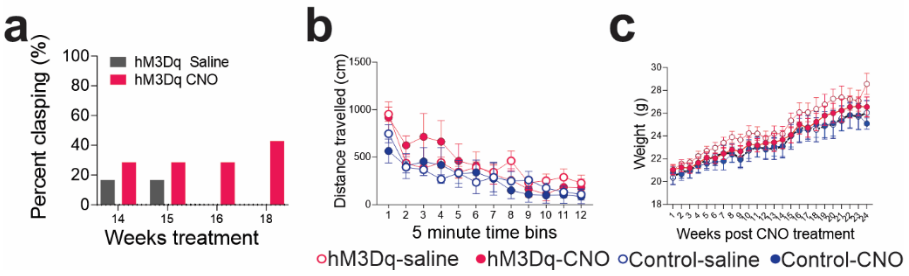


**Fig. S3. Hindlimb clasping, spontaneous locomotor activity and body weight of mouse groups.** a, Percent of hM3Dq-saline (*n*=6) and hM3Dq-CNO mice (*n*=7) clasping from 14 to 18 weeks. b, Voluntary locomotor activity in a locomotor cell did not differ between hM3Dq (*n* = 10-12 per group) and control mice (*n* = 7-8 per group) following an acute treatment with clozapine-N-oxide (CNO). c, Body weights of hM3Dq (*n* = 10-11 per group) and control mice (*n* = 7-8 per group) treated with saline or CNO mice did not differ over time. Data are expressed as mean ± SEM. Data were compared by two-way repeated measures ANOVA, $p > 0.05$.

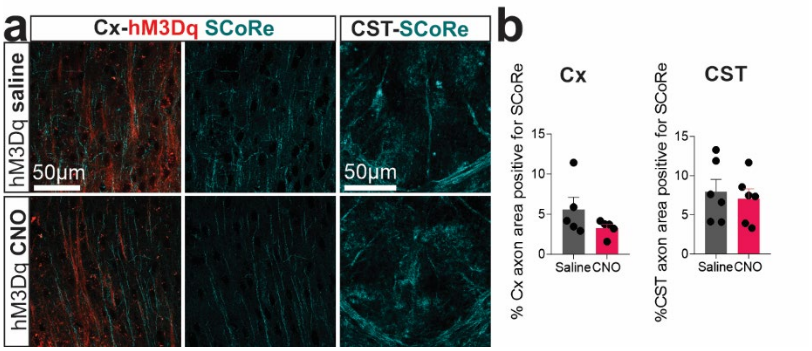


**Fig. S4. Myelin coverage is unaffected in motor cortex and corticospinal tract in hM3Dq-CNO mice.** a, Representative Spectral Confocal Reflectance (SCoRe) images of myelin signal (cyan) within the axons (red) of hM3Dq neurons, and within the dorsal funiculus of the lumbar spinal cord. b, Quantification of the myelinated area positive for SCoRe signal does not differ between hM3Dq-CNO and hM3Dq-saline mice in Layer V motor cortex (Cx), and corticospinal tract (CST) within the dorsal funiculus of lumbar spinal cord. Data are expressed as mean ± SEM. All data were compared by unpaired t-tests, $p > 0.05$, $n = 5$ per group.

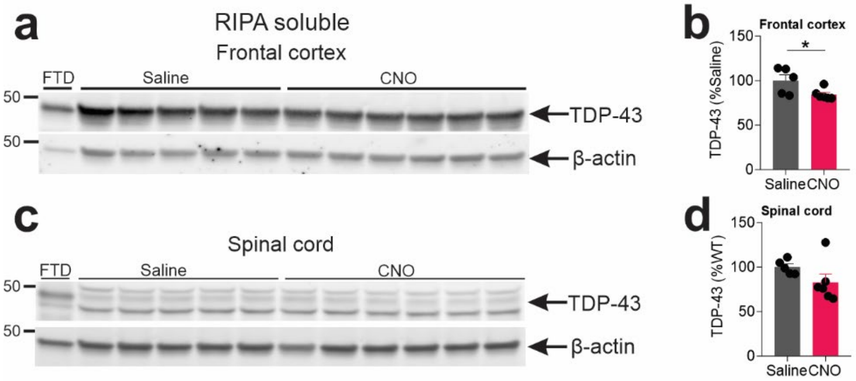


**Fig. S5. Protein levels of RIPA-soluble TDP-43 in frontal cortex are significantly reduced in hM3Dq-CNO mice.** Immunoblot analysis of RIPA-soluble TDP-43 in frontal cortex a, and c, lumbar spinal cord of hM3Dq-saline and hM3Dq-CNO mice, relative to β-actin levels. b, Quantification analysis revealing significantly lower levels of TDP-43 levels in frontal cortex of hM3Dq-CNO mice, compared to hM3Dq-saline mice. d, TDP-43 levels in lumbar spinal cord in hM3Dq-CNO mice, compared to hM3Dq-saline mice. Gel source data can be found in Supplementary Fig. 1. Data are expressed as mean ± SEM. All data were compared by unpaired t-tests, * $p < 0.05$, $n = 5\text{-}6$ per group.
